# SAPPTree: Identification of an S-Acylation Motif Drives a Novel S-Acylation Prediction Program

**DOI:** 10.64898/2026.06.30.735287

**Authors:** Alyssa S. Guo, Vu Son Luong, Andrey A. Petropavlovskiy, Anthony Dang, Andrew C. Doxey, Shaun S. Sanders, Dale D. O. Martin

**Affiliations:** Department of Biology, University of Waterloo, Waterloo, ON, Canada; Cheriton School of Computer Science, University of Waterloo, Waterloo, ON, Canada; Department of Molecular and Cellular Biology, University of Guelph, Guelph, ON, Canada; Waterloo Institute of Nanotechnology, University of Waterloo, Waterloo, ON, Canada

## Abstract

S-acylation, the reversible addition of fatty acids to proteins, has emerged as an abundant post-translational modification that drives protein localization and function. With no known consensus sequence, current prediction programs rely on machine learning algorithms that use short peptide sequences and large proteomic datasets. However, current prediction programs often suggest incorrect sites of S-acylation, leading to wasted experimental time and effort following site-directed mutagenesis and low-throughput validation experiments. Using only experimentally confirmed sites of S-acylation, we sought to identify primary sequence, secondary structure, and tertiary structure features common amongst S-acylation sites to aid in developing more robust prediction tools. In doing so, we identified an S-acylation motif including a cysteine cluster flanked by a hydrophobic stretch, and a positively charged polybasic region found within a helical stretch. These features were combined with known or AlphaFold-predicted structures and additional features including residue depth and solvent accessibility into a random forest model to generate a new and more accurate S-acylation prediction program (SAPP), named SAPPTree. All the processed datasets and complete model training pipeline are available at https://github.com/neurdyphagy-lab/palm-prediction-model, while the webserver is available at http://martintools.sci.uwaterloo.ca/.

## Introduction

As the only reversible form of protein lipidation, S-acylation has emerged as a tractable therapeutic target in a myriad of diseases(1). S-acylation involves the reversible addition of long-chain fatty acids to cysteine residues of proteins via a thioester bond. The 16-carbon saturated fatty acid palmitate is the primary lipid substrate used, and therefore S-acylation is commonly referred to as S-palmitoylation, or simply palmitoylation(2). Lipidation is mediated by a family of protein S-acyltransferases known as zinc-finger DHHC (ZDHHC) enzymes, named after their conserved Asp-His-His-Cys active site motif(3,4). The fatty acid moiety can be removed by three classes of serine hydrolases known as palmitoyl protein thioesterases (PPT), acyl protein thioesterases (APTs), and α/β hydrolase domain-containing proteins (ABHDs) that all contain an α/β hydrolase fold with a Ser–His–Asp catalytic triad and an active serine residue consensus motif of Gly–X–Ser–X–Gly(5–7). While S-acylation is best described for its role in promoting membrane binding and targeting proteins to specific locations within the cell, S-acylation is also important for mediating protein-protein interactions, protein conformation, and regulation of enzymatic activity(1,2). For example, S-acylation of active site cysteines can be used to ‘turn on/off’ enzymes, thereby acting as a potential metabolic sensor(8–10).

Not surprisingly, defects in S-acylation lead to protein mislocalization and altered function, and, as a result, are connected to many diseases, including cancers, neurodegeneration, immunological defects, and more(1,10–12). S-acylation is also key for many invading organisms, such as bacteria, parasites, and viruses, including SARS-CoV-2(13,14). As a result, S-acylation has become an attractive target in many diseases and infections(1,11). Like phosphorylation, the need to block or promote S-acylation is context-specific. Consequently, there is a great deal of interest in identifying S-acylation substrates and their sites of modification. To date, S-acyl-proteome studies have identified a plethora of potentially S-acylated proteins(10,15). However, one of the biggest impediments in characterizing these proteins is identifying the cysteine sites of S-acylation, as there is no known consensus sequence(1,2,16,17). As such, prediction programs have been developed by combining multiple data sets from various S-acylation public databases as well as literature searches(20–22). Many of these programs depend on a short number of amino acids flanking either side of the targeted cysteine and/or use large-scale high-throughput proteomic data to train the models leading to the incorporation of false-positives. Ultimately, these programs often predict false sites of S-acylation that are disproven using low throughput validation with site-directed mutagenesis(18,19).

Here, we report the identification of an S-acylation motif in which the modified cysteine is flanked by hydrophobic and polybasic domains (HPB). To identify this motif in proteins of interest, we developed a new S-acylation prediction program (SAPPTree) using a random forest model that incorporates features derived from secondary and tertiary structures. Our analyses indicate that the algorithm relies heavily on scores derived from the S-acylation HPB motif, the presence of nearby cysteine clusters, and local secondary structure. In addition, the program incorporates solvent-accessible surface area measurements for cysteine residues and neighboring amino acids calculated from protein tertiary structures. Collectively, the identification of this motif provides new insight into the mechanisms by which ZDHHC enzymes recognize and S-acylate their substrates.

## Results

### 1. Identification of an S-acylation motif

To determine if specific amino acids are preferred anywhere along the SAP(100, 100), Two Sample Logo was applied to perform a t-test for each amino acid residue at each position(23). Amino acids with a p-value less than 0.05 were displayed along SAP(100, 100) using the full amino acid alphabet (Figure 1A, Supplemental Figure 1A) and a reduced alphabet (Figure 1B, Supplemental Figure 1B). As S-acylation can occur on both C- and N-terminal sides of a transmembrane domain(24,25), a sequence or structural S-acylation motif can likely occur in both directions. Thus, the program was designed to capture these motifs regardless of orientation. When considering the full amino acid alphabet, no patterns were apparent. With the reduced alphabet, however, three distinct regions emerged as follows (Figures 1B; Supplemental Figure 1B; Table 1): a hydrophobic amino acid region (G) from position −22 to −11, −7 to −5 and at −1, a cysteine cluster region (C) surrounding the S-acylated cysteine (from position −3 to 6), and a positively charged region from position 2 to 11 and −4. Upstream of the cysteine cluster, there was an enrichment of hydrophobic amino acids including Leu, Ala, Ile, Met and Val (represented by G in black in the reduced alphabet; Supplemental Figure 1B and Figure 1B). Downstream of the cysteine cluster and the S-acylated cysteine, there was an enrichment of positively charged amino acids such as Arg and Lys (represented by K in blue in the reduced alphabet; Figures Supplemental Figure 1B and Figure 1B). When assessing secondary structure of SAP(100, 100) using Two Sample Logo’s t-test, we found a large stretch of helical amino acids enriched from position −76 to −47, −41 to +19 and at −45 (Figure 1C and Table 1). This suggests that overall, helical structures are preferred close to and surrounding the S-acylated cysteine.

**Table 1:**
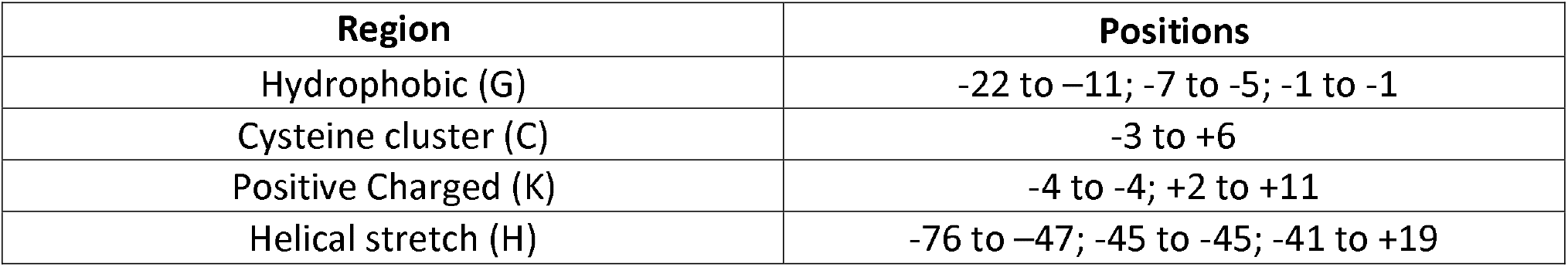
Boundaries of each primary sequence and secondary structure determinant of S-acylation identified using Two Sample Logo. The key biochemical regions identified within SAP(−100,100) and their corresponding positional coordinates relative to the S-acylated cysteine.

| Region | Positions |
| --- | --- |
| Hydrophobic (G) | -22 to -11; -7 to -5; -1 to -1 |
| Cysteine cluster (C) | -3 to +6 |
| Positive Charged (K) | -4 to -4; +2 to +11 |
| Helical stretch (H) | -76 to -47; -45 to -45; -41 to +19 |

**Figure 1:**
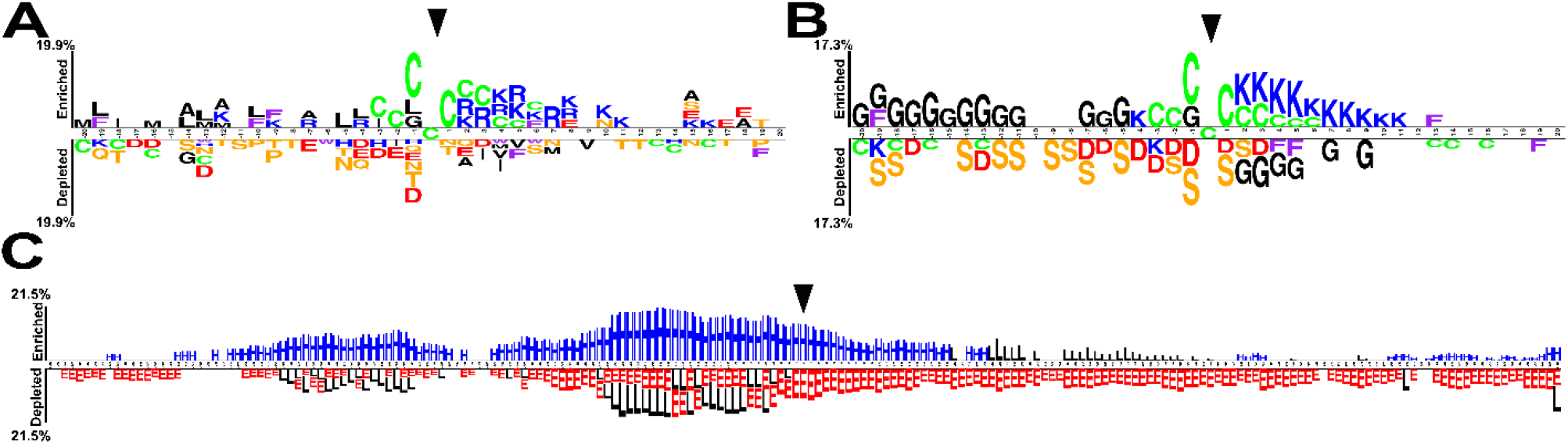
Primary sequence and secondary structure determinants of S-acylation. A) Two Sample Logo was applied to primary sequences of SAP(20, 20)s to determine amino acid preference. Amino acids with p<0.05 are shown with colour indicating the type of amino acid. B) Two Sample Logo was applied to the reduced alphabet of SAP(20, 20) with the letter codes (G=non-polar, S=polar, K=positively charged, D=negatively charged, F=aromatic, C= cysteine). C) Two Sample Logo was applied to the secondary structure of SAP(−100, 100). H = alpha-helix (blue), E = extended beta strand (red), L = loop. Black arrow indicates the S-acylated Cysteine at position 0.s

Based on these results, and understanding the mechanisms that drive S-acylation(26), we predicted that protein structure and how it relates to the position of the S-acylated cysteine would also play a role in S-acylation. Consequently, in addition to the above motif, we also included features derived from protein structures acquired from the PDB and AlphaFold(27,28), as well as solvent accessibility scores (ACC) and residue depth score. Together, these features made up the prediction program.

### 2. SAPPTree requires S-acylation Motif

Standard binary-classification evaluation metrics, including accuracy, sensitivity, specificity, and Matthew’s Correlation Coefficient (MCC), were measured in evaluating the model’s performance. Using a 70-30 train-test split, our model reached 75.67% in accuracy with sensitivity at ~74%, specificity at ~78% and MCC of 0.51 (Table 2). These scores were calculated as the average across 200 runs of training using scikit-learn’s “train_test_split” package(29).

**Table 2:** Train-test and 10-fold cross validation testing results for SAPPTree. Summary of SAPPTree’s performance when using a 70-30 Train-Test split and 10-fold cross validation.

| Category | 70-30 Train-test split | 10-fold cross validation |
| --- | --- | --- |
| Sensitivity | 73.61 % | 74.83% ± 4.80% |
| Specificity | 77.71% | 78.38% ± 4.10% |
| Accuracy | 75.67 % | 76.60% ± 3.58% |
| AUC ROC | 0.76 | 0.8546 ± 0.0272 |
| F1 | 0.7517 | 0.7615 ± 0.0371 |
| MCC | 0.51 | 0.5332 ± 0.0717 |

A 10-fold cross validation showed that our model reached similar results compared to the outcomes of the train-test split. Our model reached 76.60% in accuracy with sensitivity at ~75%, specificity at ~78% and an MCC of 0.53. The ROC of the 10-fold cross-validation and its area under the curve is similar to ROC/AUC for 4-, 6-, and 8-fold cross-validation (Figure 2).

**Figure 2:**
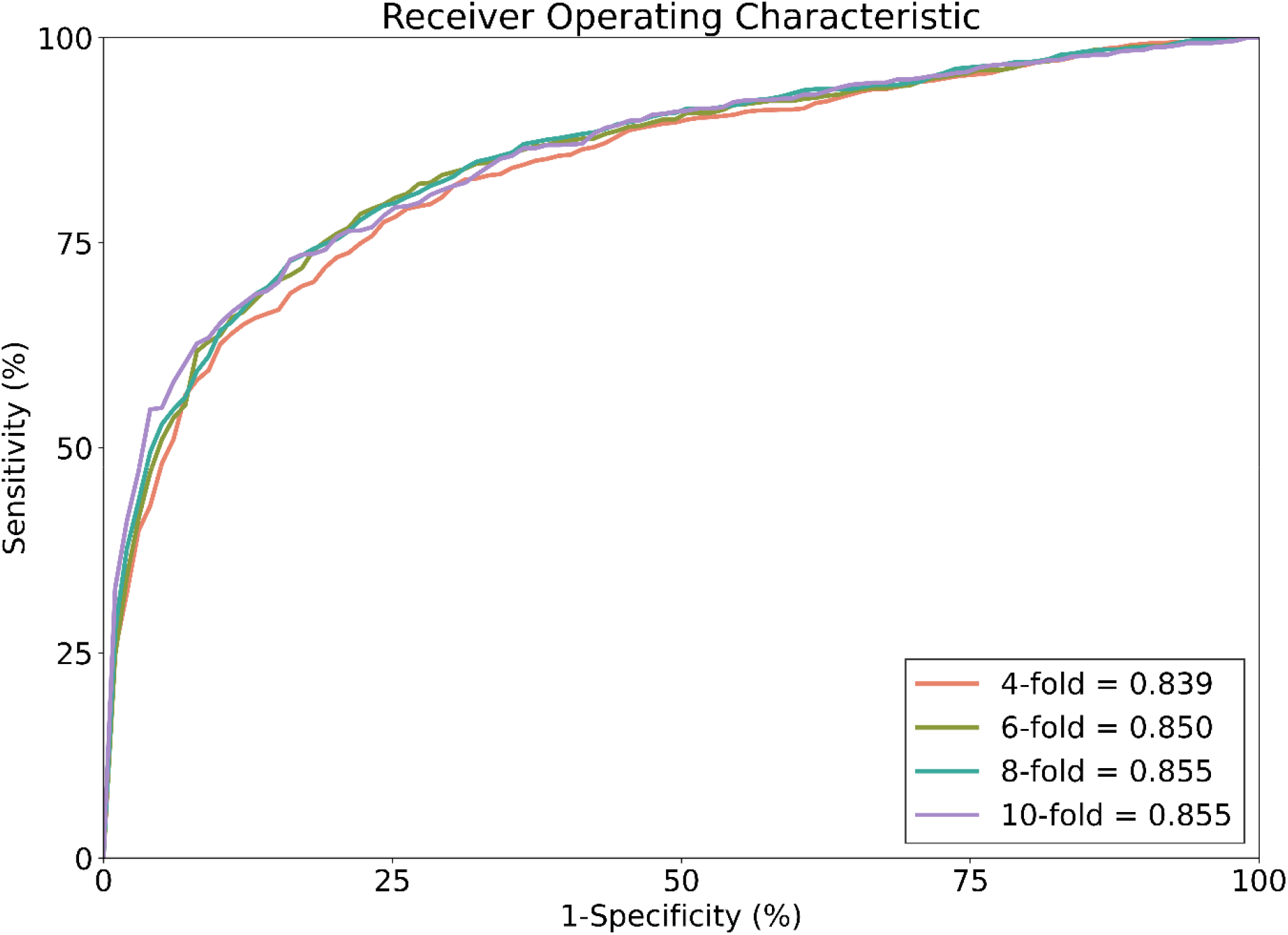
Receiver Operating Characteristic (ROC) curve across k-Fold cross-validation. The plot shows the model’s performance across 4-, 6-, 8-, and 10-fold cross validation. The curves largely overlap with the corresponding area under the curve (AUC) to be 0.839, 0.850, 0.855, and 0.855, respectively. This indicates the strong and consistent ability to distinguish between positive and negative S-acylation sites across different cross-validation settings.

In addition to model training and testing, we performed two forms of feature importance analysis to determine the extent to which each dataset feature informed the prediction model’s classification of cysteines as S-acylated or not. The feature importance internally calculated by the model during training was considered, in addition to a permutation importance analysis where performance was assessed via AUC-ROC.

The metrics indicate the proportion with which each feature is internally used in separating S-acylated versus non-S-acylated sites in the random forest model (Figure 3A). This analysis indicates that the prediction model most highly depends on the HPB domain score (comprised of the reduced alphabet features of a hydrophobic region and polybasic domain on opposite ends of the cysteine). We next performed a permutation feature analysis, which measures the importance of each feature within the random forest model using a permutation analysis rather than innate feature importance obtained during training. Under this analysis, each individual feature is randomly scrambled so that their contribution to the model is no longer relevant. Then, the decrease in performance caused by losing this feature is measured by calculating the average drop in ROC-AUC across 10 independent runs. The highest decrease in performance was observed when the HPB domain score and cysteine cluster score (which measures the presence of multiple cysteines) were removed from the model (Figure 3B). Coupled with the internal model feature importance (Figure 3A), alongside the model performance (Table 2), these findings indicate that the model relies heavily on the HPB and helical domains score to predict S-acylation, with a high degree of accuracy. Similarly, feature importance analysis performed on the entire dataset showed that the model was highly dependent on scores generated from the HPB domain (Figure 3C). In addition, permutation feature analysis revealed a significant drop in model performance when motif-related features were removed (Figure 3D). For the remaining features that are related to solvent accessibility and residue depth, the feature importance analysis indicated that the model still relied substantially on them (Figure 3C). Although removing these features individually did not significantly decrease model performance in this case (Figure 3D), the feature importance analysis (Figure 3C) indicated they still contributed to the model and thus were left in the final model.

**Figure 3:**
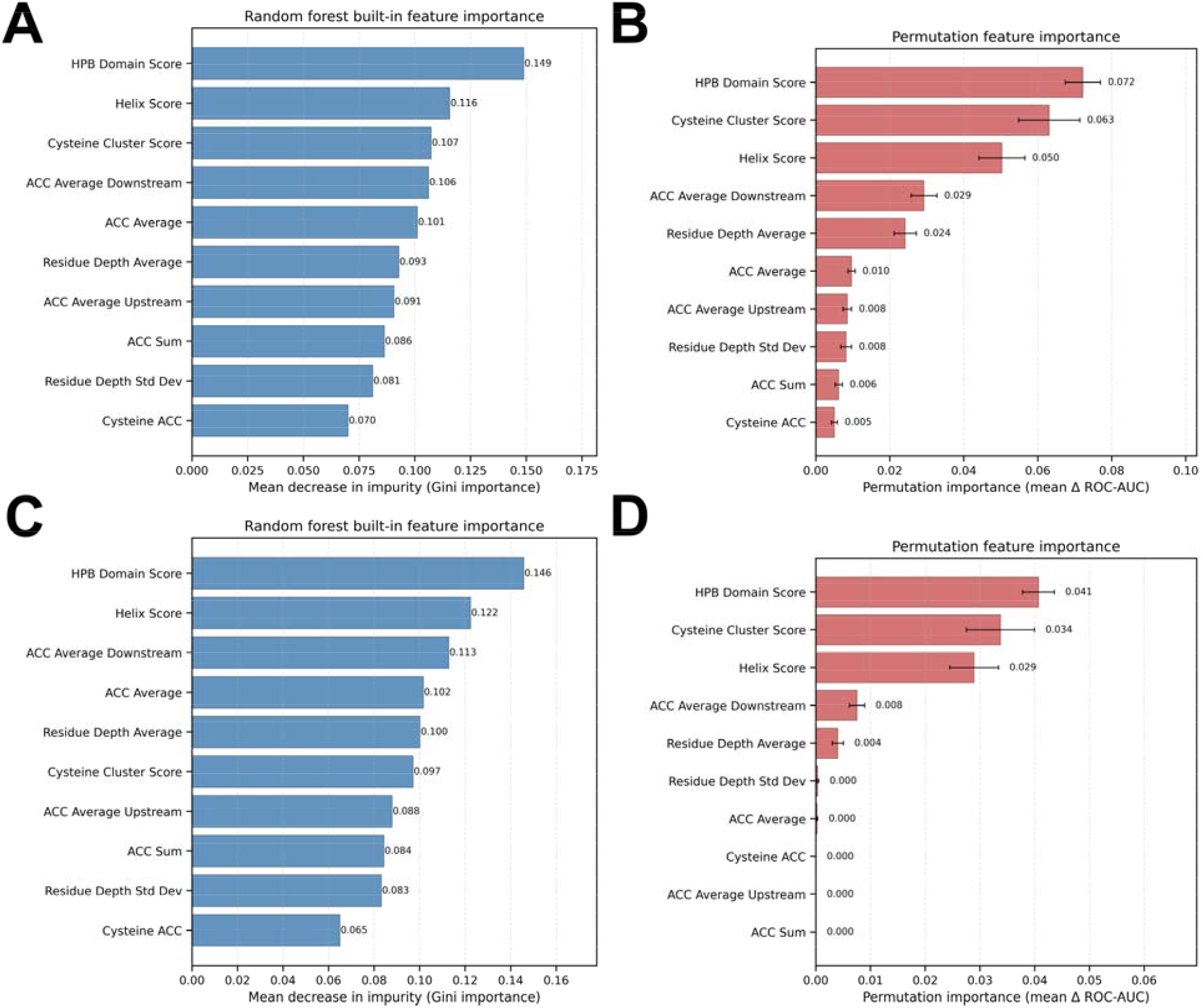
SAPPTree input feature importance. A) Relative internal feature importance to the random forest model obtained after training using 70% of sites. The scores sum up to 1, with a higher score indicating that a feature is more strongly relied on. B) Average change in model performance observed from 10 runs of permutation feature analysis using 70% of sites for training. C) and D) represent the same results from A) and B), respectively, using all positive sites. Error bars denote standard deviation.

### 3. SAPPTree outperforms common S-acylation prediction program

We next sought to compare SAPPTree against a previously released S-acylation predictor program. As GPS-Palm was developed from CSS-Palm 1-4(20,22), which are among the most highly used programs, and linked to all proteins included in Swisspalm(15), this was chosen as the benchmark to evaluate SAPPTree. Comparing the positive datasets used to train GPS-Palm and SAPPTree, both SAPPTree and GPS-Palm shared 397 S-acylated sites. SAPPTree has 366 unique positive sites while GPS-Palm has 2701 unique positive sites (Figure 4). This large number in GPS-Palm primarily comes from the use of large proteomic datasets(22). To directly compare performance of GPS-Palm and SAPPTree, a 70-30 train-test split was applied to the SAPPTree data, with all 397 shared sites included in the training dataset for SAPPTree, and the remaining 366 unique sites split between the training and testing datasets to achieve a 70-30 split (Figure 4). As GPS-Palm’s negative sites used for training are not publicly available, SAPPTree’s negative dataset was randomly sampled into a 70-30 train-test split.

**Figure 4:**
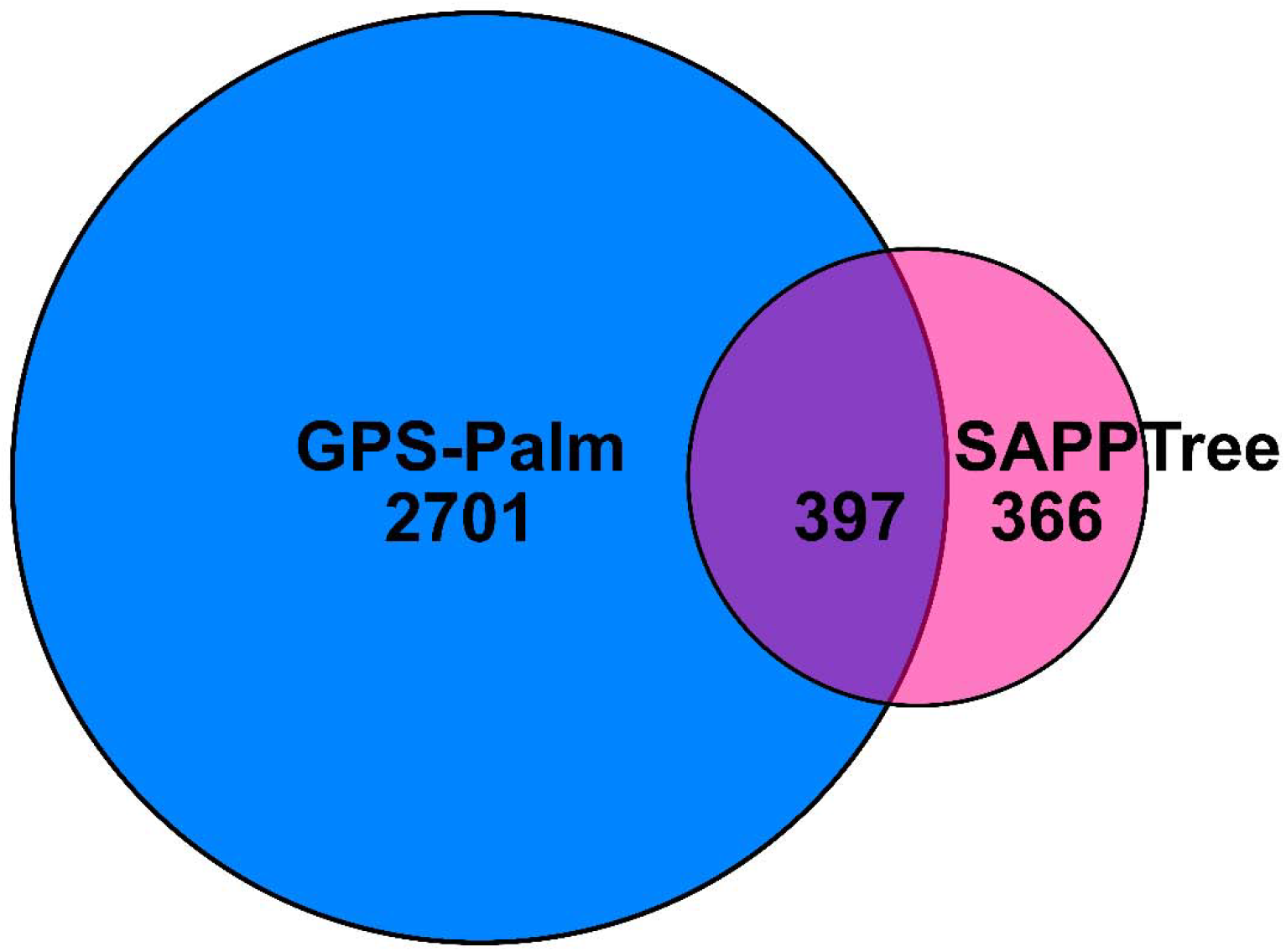
Comparison of positive data sets used in SAPPTree and GPS-Palm. 397 S-acylation sites were common to both datasets. GPS-Palm contained a larger number of unique positive S-acylation sites including 2,701 sites that are not present in SAPPTree. In contrast, SAPPTree included 366 unique positive S-acylation sites not in GPS-Palm. Overall, the comparison indicates limited overlap between the two datasets, with GPS-Palm contributing the larger dataset-specific positive set.

Overall, SAPPTree was more accurate than GPS-Palm (72.93% vs ~56%), with a higher sensitivity, and specificity (Table 3). Overall, this means SAPPTree more often predicts S-acylation sites (sensitivity) with fewer false positives (specificity). In addition, the MCC was nearly three times higher for SAPPTree than for GPS-Palm (Table 3), indicating this is a robust prediction program. A final comparison using all sites used in SAPPTree (Table 4), including the 397 sites shared with GPS-Palm, shows SAPPTree is better all around at predicting sites with higher sensitivity and accuracy.

**Table 3:** Comparison of GPS-Palm and SAPPTree when testing against 229 unique positive S-acylation. SAPPTree outperformed GPS-Palm in sensitivity, specificity, accuracy, and MCC on the independent positive-site test set (for 70/30 train/test split in SAPPTree). GPS-Palm sets thresholds of low (0.6484), medium (0.7766), and high (0.8920) based on specificity.

| Program | GPS-Palm |  |  | SAPPTree |
| --- | --- | --- | --- | --- |
| Threshold | low | med | high | Average |
| Sensitivity | 60.26% | 40.61% | 28.38% | 67.86% ± 1.28% |
| Specificity | 53.28% | 68.56% | 85.15% | 77.99% ± 0.994% |
| Accuracy | 56.77% | 54.59% | 56.77% | 72.93% ± 0.84% |
| MCC | 0.14 | 0.10 | 0.16 | 0.4609 ± 0.0168 |

**Table 4:** When testing against the entire positive SAPPTree dataset (763 sites): SAPPTree provided stronger, more balanced performance overall, with higher sensitivity, specificity, accuracy, and MCC than GPS-Palm.

| **Program** | **GPS palm** |  |  | **SAPPTree** |
| --- | --- | --- | --- | --- |
| **Threshold** | **low** | **med** | **high** | **Average** |
| **Sensitivity** | 69.72% | 53.47% | 34.99% | 74.83% ± 4.80% |
| **Specificity** | 50.98% | 67.37% | 85.45% | 78.38% ± 4.10% |
| **Accuracy** | 60.35% | 60.42% | 60.22% | 76.60% ± 3.58% |
| **MCC** | 0.21 | 0.21 | 0.24 | 0.5332 ± 0.0717 |

## Discussion

Over the past few decades, protein S-acylation has been recognized as an important post-translational modification that regulates vital signaling pathways(1,7,12,25,30). S-acylation dynamically modulates many protein properties, including stability, localization, protein-protein interactions, and function, but is most commonly known for its role in directing protein localization. As a result, S-acylation regulates cellular physiology, and dysregulated S-acylation has been associated with multiple human diseases.

Experimentally validating whether a protein is S-acylated typically requires selecting candidate S-acylated cysteine residues, generating cysteine to alanine or serine mutations, and performing S-acylation assays (e.g. acyl-biotin exchange or metabolic labeling and click chemistry) following exogenous expression of the mutant proteins in cells. Depending on the protein, screening each cysteine using this approach can be time-consuming. Thus, accurate prediction tools are essential for prioritizing candidate sites. To date, multiple S-acylation predictors have been developed(20–22) and CSS-Palm is a widely used program for S-acylation site prediction that has been incorporated into the online database of all S-acylated proteins, SwissPalm(15). Ultimately, the creators of CSS-Palm replaced the program with GPS-Palm(22). Since then, SwissPalm has had two new releases, and the number of validated S-acylation sites in *Homo sapiens, Mus musculus*, and *Rattus norvegicus* has increased from 1,064 sites in 2019 to 1,204 sites in 2022 and 1,303 sites in 2024(15). Therefore, an updated and improved predictor for S-acylation sites is needed.

Recently developed S-acylation site prediction programs have begun using high-throughput datasets that incorporate the sites obtained by using mass spectrometry-based protein identification strategies(22,31–33). Although such methods are useful for identifying S-acylated sites across the proteome, they often contain many false-positive and false-negative sites, which can significantly reduce model performance(22). Consistent with this limitation, when we tested our low-throughput dataset of 763 experimentally verified sites using GPS-Palm, the model performance was substantially lower than previously reported.

To improve data quality, we only used S-acylation sites verified by low-throughput approaches to train the model. In addition, since Ning *et al*.(22) previously found significant differences among N-terminal, C-terminal, and internal S-acylated cysteines, we constructed the negative dataset with equivalent numbers of N-terminal, C-terminal, and internal cysteines to better reflect the distribution of S-acylation sites. Furthermore, solvent accessibility and residue depth were calculated based on the tertiary structure of the protein rather than the primary peptide sequence. Together, these improvements led to a significant increase in the performance of the SAPPTree model compared with earlier prediction programs.

The reduced-alphabet and secondary-structure Two Sample Logo results suggest that the S-acylation motif is determined by three-dimensional structural features in addition to the primary amino acid sequence, rather than primary sequence alone. The S-acylated cysteine cluster is flanked by a 22-amino-acid hydrophobic region and an 11-amino-acid polybasic region (Figure 1B; Supplementary Figure 1B). In addition, the Two Sample Logo analysis revealed a stretch of helical secondary structure across SAP(100, 100) (Figure 1C). However, examination of individual SAP(100, 100) secondary structures showed that these helices are typically 10–30 amino acids long and can occur either upstream or downstream of the S-acylated cysteine.

Furthermore, the helices within SAP(100, 100) and HPB domain may help anchor the sequence to the cell membrane, where the transferase active site is located. The current model of S-acylation involves two catalytic steps(26,34). First, the cysteine residue within the DHHC catalytic site of the S-acylation transferase undergoes auto-acylation. The acyl group is then transferred from the DHHC cysteine to the central cysteine of the substrate protein within SAP(100, 100). Structurally, the catalytic site of the transferase is positioned between two of the transmembrane domains in close proximity to the membrane(26). Based on these observations, we hypothesize that the upstream hydrophobic region and downstream polybasic region flanking the S-acylated cysteine help position the central cysteine so that it aligns properly with the DHHC catalytic site during the S-acylation reaction. Alternatively, or additionally, the polybasic region may interact with negatively charged phospholipids on the cell membrane. Furthermore, the enriched helices within SAP(100, 100) and HPB domain may help anchor the sequence to the cell membrane, where the transferase active site is located.

As more protein structures are resolved and as AlphaFold evolves, along with the confirmation of more S-acylation sites, our program is set to become more accurate over time. The identification of the S-acylation motif containing the HPB domain and helical stretch will contribute to our understanding of how S-acylation is mediated by the ZDHHC enzymes.

Ultimately, SAPPTree serves as a valuable tool for the S-acylation community as it overcomes many accuracy limitations of the previous prediction programs and allows researchers to save time and accelerate discovery of new S-acylation sites.

## Data and code availability

The processed datasets used to train SAPPTree, pre-trained model, and code files required to reproduce the training and analysis pipeline are all available at https://github.com/neurdyphagy-lab/palm-prediction-model. The web-server hosting SAPPTree for users to make S-acylation predictions is available at http://martintools.sci.uwaterloo.ca/.

## Materials and Methods

### 1. Benchmark Dataset Generation

We obtained data of experimentally verified S-acylation sites from *Homo sapiens, Mus musculus*, and *Rattus norvegicus* in the 5 th release of SwissPalm(15). These S-acylation sites were confirmed by site-directed mutagenesis followed by low-throughput S-acylation identification methods (e.g. acyl-biotin exchange, metabolic labeling followed by click chemistry). The primary protein sequences were obtained from the SwissPalm-linked Uniprot database(35). Cluster Database at High Identity with Tolerance (CD-HIT)(36) was used to remove protein sequences that were >50% in identity to avoid model overfitting. Where applicable, proteins excluded by CD-HIT based on similarity but known to be distinct (e.g. Fyn, Ras, etc.) were manually added back to the database (Supplemental Table 1). By mapping the sites on those protein sequences, we collected 763 verified S-acylated sites from 397 proteins (Supplemental Table 1).

As previously described(22), the S-acylation peptide (SAP(*m, n*)) has a center cysteine residue with *m* residues upstream and *n* residues downstream. Approximately 20% of all sites were within 100 residues of the N- or C-terminus, leading to values of *m* or *n* less than 100. For these sites, a gap “-” was added at the corresponding N- or C-terminus to normalize the length to 201. Consequently, the parameter was set to *m* = *n* = 100. SAP(100, 100)s derived from known S-acylated cysteines were used for the positive dataset while the remaining SAP(100, 100)s from other non-S-acylated cysteines within the same proteins were randomly sampled to be used for the negative dataset. The number of positive cysteines at the N- and C-termini were also considered for the random sampling of the negative dataset, such that the negative dataset had the same number of C-terminal, N-terminal, and internal cysteines. To summarize, we constructed a dataset containing 763 positive sites and 763 negative sites from 397 proteins. Both datasets contained an equal number of N-terminal, C-terminal, and internal center cysteines.

### 2. Position Weight Matrix Generation

For a group of peptides, the probabilities of each position were considered independent and the frequencies of a specific character (an amino acid or a letter in reduced alphabet) at a particular position was calculated. The sum of frequencies at a particular position was assigned 1 and these probabilities were then converted to weights using the following formula, where background is the average probability of a character in a negative dataset (37):

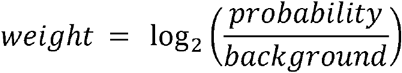

Position weight matrix was calculated from the reduced alphabet and secondary structure SAP(100, 100).

### 3. Feature Extraction

#### a) Two Sample Logo

Two Sample Logo(23) was used to visualize the statistical differences between the positive and negative SAP(100, 100)s at each position. The symbols at specific position were shown as enriched or depleted if it passed the statistical t-test with p-value < 0.05. The resulting graphic was used to determine the location on SAP(100, 100)s to calculate derivative features.

#### b) Reduced alphabet

To enable the model to extract more generalizable biochemical features, all SAP(100, 100)s were converted to a reduced alphabet. Amino acids were divided into cysteines (C) and 5 major groups: positively charged (K, R, H) represented by the letter K, negatively charged (D, E) represented by the letter D, polar (S, T, N, Q, P) represented by the letter S, non-polar (G, A, V, L, M, I) represented by the letter G, and aromatic amino acids (F, W, Y) represented by the letter F. By applying a position weight matrix to the reduced alphabet SAP(100, 100)s, the hydrophobic scores, cysteine cluster scores, and positively charged amino acid scores were calculated as the sum of the score of the associated reduced alphabet letter. The hydrophobic and positively charged amino acid scores were combined to produce the hydrophobic polybasic (HPB) domain score. Additionally, each sequence was scored in the forward or reverse direction (i.e. a flipped HPB motif) and the highest of the two was used.

#### c) Secondary structure

Experimentally verified protein structures of each S-acylated protein in protein data bank (PDB) file format were downloaded from Research Collaboratory for Structural Bioinformatics (RCSB) PDB(27). In the case that no verified structure was available or verified structures had a coverage of less than 90 percent, the predicted protein structure was downloaded from the AlphaFold database(28). The secondary structures of SAP(100, 100) were then extracted from these PDB files using the Dictionary of Protein Secondary Structure (DSSP) program(38,39). Notably, there were some edge cases in which AlphaFold does not have a predicted structure available, such as non-human proteins exceeding 1280 amino acids(28). Therefore, for some usages of the program where neither a verified structure nor an AlphaFold entry is available, the program may currently fail. Two Sample Logo(23) was applied to the extracted secondary sequences to identify the location of the alpha-helix enrichment in SAP(100, 100). As described above for the reduced alphabet, the position weight matrix was then used to calculate the helical score in SAP(100,100). Moreover, two sets of helical scores for each secondary structure were generated as follows with the highest score used for the final prediction: one scoring the presence of an upstream helical region and the other scoring the presence of a downstream helical region.

#### d) Residue depth scores

Using the Biopython package(40), MSMS program(41), and collected protein structures (experimental or predicted by AlphaFold), the residue depth scores of each amino acid in a SAP(100, 100) were calculated. Using these position-based scores, the average and standard deviation of residue depth in a SAP(100, 100) were calculated. If the protein structure was too large or had a complicated geometric complexity that caused the MSMS program(42) to fail, it was set at 0.

#### e) Solvent accessibility scores

Using PDB files and the Dictionary of Protein Secondary Structure (DSSP) DSSP program, the solvent accessibility (ACC) scores of each amino acid in a SAP(100, 100) were calculated(38,39). Using the individual ACC score of each amino acid, the sum and average ACC score were calculated. Due to approximately 20% of the SAP(100, 100)s being positioned within C- and N-termini sequences, the sum and the average of ACC scores were treated as different features. Furthermore, the ACC score of the center cysteine as well as the averages of ACC scores of the upstream and downstream region were also used as input features.

### 4. Model construction

#### a) Machine learning model selection

By applying the features extracted from SAP(100, 100) into a random forest model, a prediction model for S-acylation sites was created (Supplementary Table 1). A random forest model is a machine learning model comprised of multiple decision trees, whose predictions are individually evaluated on the dataset before taking the majority vote as the final result(43). Each decision tree is comprised of different branches that split off depending on a random subset of data benchmarks, leading to a final outcome after a series of decisions are made. As the random forest model was implemented using scikit-learn’s built-in packages(29), the comprised dataset was used as input, from which the best set of decision trees was automatically chosen to create the final random forest model (Supplementary Table 1). This model was chosen for this purpose because the ability of a random forest model to average predictions across multiple decision trees prevents overfitting, allows for feature importance analyses, and is robust to outliers and missing data(43).

#### b) Hyperparameter Selection Process

The final hyperparameters (Table 5) inputted into the random forest model package were chosen by performing a two-step optimization process. In the first step, a GridSearchCV(29) was performed using the following benchmark values outlined in Supplementary Table 2. At this stage, the parameters that typically affect random forest model performance the most were prioritized. A GridSearchCV trains the model on every single possible combination of hyperparameters as outlined in the grid, then evaluates their performance to output the best hyperparameters(29). Here, the Area Under the Curve (AUC) of the Receiver Operating Characteristic (ROC) Curve was used to evaluate performance(29). This metric was chosen as it accounts for both the sensitivity and specificity of the model. In the second step, a GridSearchCV was performed across a finer grid constructed using the hyperparameters from the first step that yielded the best scores, alongside the directly neighboring hyperparameter values. For instance, if “n_estimators=100” yielded the highest score in the first iteration, the second iteration would include “n_estimators=50”, “n_estimators=100”, and “n_estimators=200”. In addition, the hyperparameters “max_features”, “class_weight”, “criterion”, and “bootstrap” were included as part of the grid. Following this step of evaluation, the highest scoring hyperparameters using the AUC-ROC metric were ultimately chosen as inputs to the model.

**Table 5:** Hyperparameter inputs to scikit-learn’s random forest Classifier package. The hyperparameter values used in the final random forest model package to construct the model.

| Hyperparameters | Value |
| --- | --- |
| N_estimators | 200 |
| Min_samples_split | 2 |
| Min_samples_leaf | 1 |
| Max_Features | 'sqrt' |
| Max_depth | 20 |
| Bootstrap | False |
| Random_state | 42 |

#### c) Feature Importance Analysis

Feature importance analysis was also run over the predictive model, using both the built-in feature importances made available through scikit-learn’s interface, and a permutation importance analysis(29). Scikit-learn’s “feature_importances” is built into its random forest model package, which is calculated during training and measures the relative degree to which each individual feature in the dataset contributes to differentiating between positively versus negatively labelled samples (i.e. S-acylated versus non-S-acylated)(29). The permutation importance analysis was run using scikit-learn’s “permutation_importance” method, which similarly measures feature importance, but by measuring the degree to which the model’s performance decreases when individual features are scrambled so that they are no longer meaningful inputs(29).

### 5. Web server

To make SAPPTree available for usage, we developed a webserver that, upon receiving a Uniprot ID as input, runs the same calculations and analyses detailed in the methods for all cysteines to predict whether they are S-acylated. The webserver utilizes the same sequence of analysis detailed in the methods to arrive at a prediction (Figure 5) and can process one protein at a time. The SAPPTree web server is available for usage at http://martintools.sci.uwaterloo.ca/.

**Figure 5:**
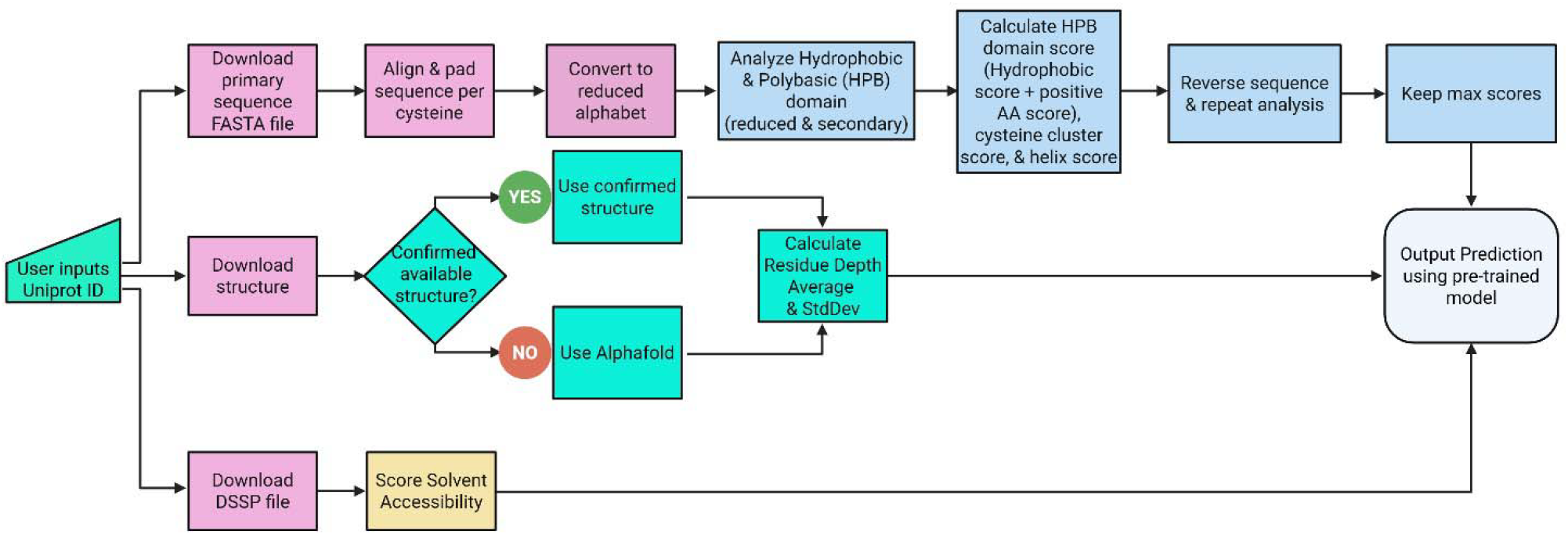
Program architecture from user input to prediction output. Arrows show the flow of data between various components, which implement different steps of analysis described in the methods. The same steps of analysis were performed to derive the dataset used to train and test the model.

### 6. Performance evaluation

The prediction model was evaluated using both train-test split and 10-fold cross validation. Under these evaluations, standard evaluation metrics, including sensitivity (Sn), specificity (Sp), accuracy (Ac), F1 score, and Matthews’s Correlation Coefficient (MCC), were calculated using scikit-learn’s “train_test_split” package(29). F1 score represents the harmonic mean between the precision (out of all predicted positive, how many are truly positive) and recall (out of all actual true positive, how much the model predicts correctly). The other evaluation metrics are defined below. For the train-test split, the comprehensive dataset was split so that 70% (966 samples) was used in training the model, and the remaining 30% (414 samples) was used to evaluate the model performance. This train-test split was run 200 times, and the average Sn, Sp, Ac, F1, and MCC across all runs were measured to evaluate the model.

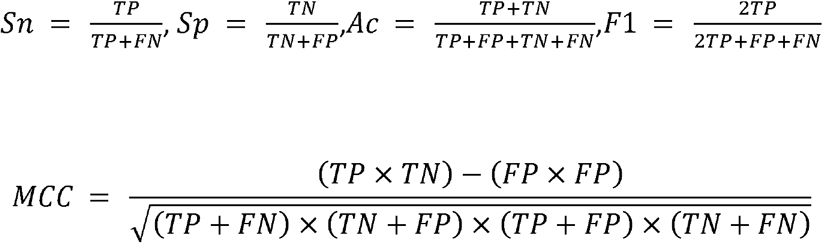

10-fold cross validation on the random forest model was performed by using the “cross_val_score” method within scikit-learn’s “model_selection” package(29). The ROC curve was also generated for the model by using this package from scikit-learn.

## Acknowledgements

This work was supported by the Natural Sciences and Engineering Research Council of Canada (grant numbers RGPIN-2021-02547 to SSS, RGPIN-2019-04617 to DDOM, and RGPIN-2025-04130 to AD). Anthony Dang held an NSERC Undergraduate Student Research Award. This research was conducted on the Dish with One Spoon territory of the Mississaugas of the Credit of the Hodinöhsö:ni and Anishinaabe peoples, land that is home to past, present, and future First Nations, Inuit, and Métis peoples. A special thank you to Luc Berthiaume who can very accurately predict sites of S-acylation by eye, but whom we could not cite in papers when trying to select potential sites to target for site-directed mutagenesis.

