## Supplemental Figure 1 for "SAPPTree: Identification of an S-Acylation Motif Drives a Novel S-Acylation Prediction Program"

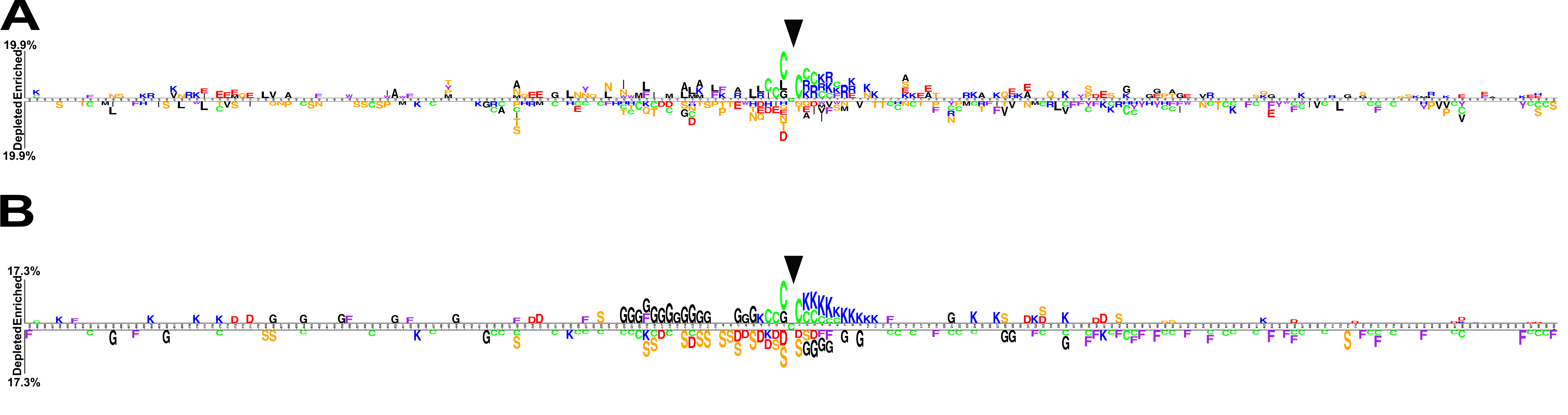

**Supplementary Figure 1**: **Primary sequence and secondary structure determinants of S-acylation.** **A**) Two Sample Logo was applied to primary sequences of SAP(100, 100)s to determine amino acid preference. Amino acids with p<0.05 are shown with colour indicating the type of amino acid. **B**) Two Sample Logo was applied to the reduced alphabet of SAP(100, 100) with the letter codes (G=non-polar, S=polar, K=positively charged, D=negatively charged, F=aromatic, C= cysteine).
