## Supplemental Table 2 for "SAPPTree: Identification of an S-Acylation Motif Drives a Novel S-Acylation Prediction Program"

**Supplementary Table 2: Candidate hyperparameter inputs to scikit-learn's GridSearchCV package.** The hyperparameter values used to create test combinations for hyperparameter tuning.

| **Hyperparameters** | **Candidate values** |
| --- | --- |
| N_estimators | [50, 100, 200, 300] |
| Min_samples_split | [2, 5, 10] |
| Min_samples_leaf | [1, 2, 4] |
| Max_Features | ["sqrt", "log2", None] |
| Max_depth | [5, 10, 20, 30] |
| Bootstrap | [True, False] |
